# Characterization of spatial dynamics of fMRI data in white matter using diffusion-informed white matter harmonics

**DOI:** 10.1101/2020.10.28.359125

**Authors:** Hamid Behjat, Iman Aganj, David Abramian, Anders Eklund, Carl-Fredrik Westin

## Abstract

In this work, we leverage the Laplacian eigenbasis of voxelwise white matter (WM) graphs derived from diffusion-weighted MRI data, dubbed *WM harmonics*, to characterize the spatial structure of WM fMRI data. Our motivation for such a characterization is based on studies that show WM fMRI data exhibit a spatial correlational anisotropy that coincides with underlying fiber patterns. By quantifying the energy content of WM fMRI data associated with subsets of WM harmonics across multiple spectral bands, we show that the data exhibits notable subtle spatial modulations under functional load that are not manifested during rest. WM harmonics provide a novel means to study the spatial dynamics of WM fMRI data, in such way that the analysis is informed by the underlying anatomical structure.

## 1. INTRODUCTION

Despite past controversies in relation to the source of the blood-oxygen-level-dependent (BOLD) signal in white matter (WM) [1], reports of fMRI activation [2, 3] and functional connectivity [4, 5] in WM continue to increase. The BOLD signal in WM has been shown to exhibit a spatial correlational anisotropy that coincides with underlying fiber patterns [6], which is manifested both at rest and under varying functional loads [7]. In addition, it has been shown that the dynamics in WM are concomitant with those observed in cortical regions connected through fiber bundles [7, 8]. The unique spatial structure of the BOLD signal in WM warrants a revisiting of conventional methods used for spatial processing of fMRI data, in particular, the use of isotropic (often Gaussian) smoothing kernels to preprocess the data [9]. Given the body of recent reports on the anisotropic spatial structure of WM fMRI data, the implicit assumption on the isotropy of the BOLD contrast that justifies use of isotropic filters may not hold in WM.

Recent work implementing spatial smoothing on diffusion-informed WM graphs has shown the benefit of using anisotropic filters that adapt to the underlying diffusion structure in WM [10, 11]. Such WM graphs have also been found beneficial in showing the collective mediation of WM pathways based on functional activity in gray matter [12]. The present work builds on the benefits of subject-specific, voxel-wise WM graphs, showing that their Laplacian eigenbasis, dubbed *WM harmonics*, can provide a novel means for quantifying the spatial structure in WM fMRI data. In particular, using principles from the recently emerged field of graph signal processing (GSP) [13, 14], we decompose WM fMRI data using a systems of spectral kernels that covers fine-scale as well as coarse-scale bands across the spectrum. We then quantify the spectral energy (SE) content of WM fMRI data at different spectral bands across the graph spectra, and show that under functional load, spatial patterns correspond to more spatially varying WM harmonics that encode subtle anisotropic spatial patterns confined by the underlying diffusion structure arise. In contrast, we show that the observed temporal modulation under functional load is not present during rest.

## 2. METHODS

### 2.1. Dataset

We studied data from the “100 Unrelated Subjects” group (54% female, mean age = 29.11± 3.67, age range = 22-36) of the Human Connectome Project (HCP) dataset [15], which we denote as HCP100. Five of the subjects were excluded due to incomplete WM coverage of the diffusion MRI data, leaving a total of 95 subjects. We used the minimally pre-processed structural MRI, diffusion MRI, task fMRI and resting-state fMRI data of all subjects; the functional data were resampled to the resolution of the diffusion data, 1.25 mm isotropic. For the task data, we studied the “Social” task, which consists of two experimental conditions: *mental* and *random*. During the mental-condition blocks, participants were presented with short video clips (20 seconds) of objects (circles, squares, triangles) that interacted in some way, whereas during the random-condition blocks the objects moved randomly on the screen. The paradigm consisted of 3 mental-condition and 2 random-condition trials. The proposed analysis scheme relies on accurate co-registration between structural, diffusion, and functional data, which is meticulously performed on the preprocessed HCP data. A thorough description of the image acquisition parameters and preprocessing steps can be found in [16].

### 2.2. Graph signal processing fundamentals

Consider an undirected, weighted graph consisting of *N* vertices in which the edges and their weights are given by an *N* × *N* adjacency matrix, with elements *a_i,j_* > 0 if an edge connects vertices *i* and *j*, and *a_i,j_* = 0 otherwise. The graph’s normalized Laplacian matrix **L** is defined as 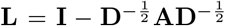, where **D** denotes the graph’s degree matrix, with its diagonal elements given as *d_i_,_i_ = ∑_j_ a_i,j_*, and **I** denotes the identity matrix. The eigendecomposition of **L** gives **L** = **UΛU\***, where **Λ** is a diagonal matrix with the eigenvalues 0 = λ_1_ ≤ λ_2_ … ≤ λ_*N*_ ≤ 2 on its diagonal— which define the graph’s *spectrum* [17], and **U** is a matrix of eigenvectors, wherein each column **u**_*k*_ is the eigenvector associated to λ*_k_*. Hereon, we refer to the eigenvectors as the *eigenmodes* as conventionally used by the neuroimaging community. The eigenbasis of **L** entails a notion of spatial variability. That is, a given eigenmode u_*k*_ is linked to its associated eigenvalue as **u**_*k*_^*T*^ **Lu**_*k*_ = λ_*k*_, which shows that *λ_k_* is a measure of total variability of **u**_*k*_. As such, eigenmodes associated to larger eigenvalues entail a greater extent of spatial variability than those associated to smaller eigenvalues.

Let **f** ∈ ℝ^*N*^ denote a graph signal, where **f**[i] is the value of the signal at vertex *i*, and let 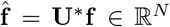 denote its spectral representation, which satisfies the Parseval energy conservation relation, i.e., 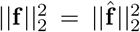.Given a continuous kernel defined on the spectral range of the graph, denoted *k*(·): [0, λ_*N*_] → ℝ, a graph signal f can be filtered using *k*(·), denoted k(**L**)**f** ∈ ℝ^*N*^, as [13]

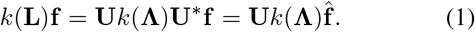

To avoid explicit computation of **Λ** and **U**, i.e., diagonalization of **L**, filtering can be alternatively done in a computationally efficient way using a polynomial approximation of *k*(·), denoted *k*_*p*_(·): [0, λ_*N*_] → ℝ, as

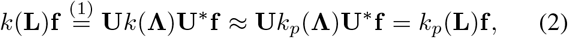

that is, computing a set of matrix operations on **L** and applying the resulting matrix to **f**.

### 2.3. Diffusion-informed WM graph design

To characterize the underlying *domain* of WM fMRI data, we leveraged diffusion-weighted MRI data to construct voxel-resolution graphs based on the method proposed in [11]. In particular, for each subject, and each hemisphere, we constructed a graph, wherein each WM voxel is represented as a graph vertex, and the relation between neighboring voxels is defined based the extent of coherence between their associated diffusion ODFs: two vertices whose associated voxels are adjacent are connected through an edge with a high weight if their associated ODFs are well aligned with the edge connecting them, and vice versa. For further details on the design we refer to [11]. We refer to the Laplacian eigenmodes of the resulting WM graphs as *WM harmonics.*

### 2.4. Spectral characterization of WM fMRI data

We represent fMRI volumes as graph signals. In particular, given a 4-D fMRI time series dataset of a given subject s, we represented the fMRI volume associated to each time instance *t* as a graph signal, denoted **f**_*s*,*t*_ ∈ ℝ^*N*^, through extracting functional values at voxels corresponding to the graph vertices, i.e., voxels within WM. The signal was then de-meaned and normalized as:

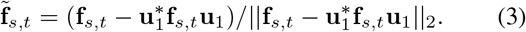

Given the sheer size of voxel-wise WM graphs, diagonalizing the associated **L** is impractical, and therefore, computation of the signal’s spectral representation, i.e., 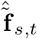, is infeasible. Instead, to obtain an overall estimate of the spectral energy (SE) content of fMRI data on WM graphs at fine-scale spectral bands (FSB), we decomposed the signals using the system of 57 spectral kernels presented in [18], see Fig. 1(a), denoted 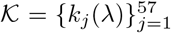, which form a Parseval frame, i.e., ∀λ ∈ [0, λ_*N*_], 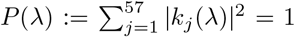, a property that ensures energy conservation between the vertex and spectral representations of the signal, i.e., 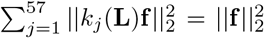 [19]. Specifically, we leveraged tailored Chebyshev polynomial approximations of 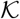, with kernel-specific polynomial orders of mean 300 ± 200, to filter signals as in (2).

**Fig. 1:**
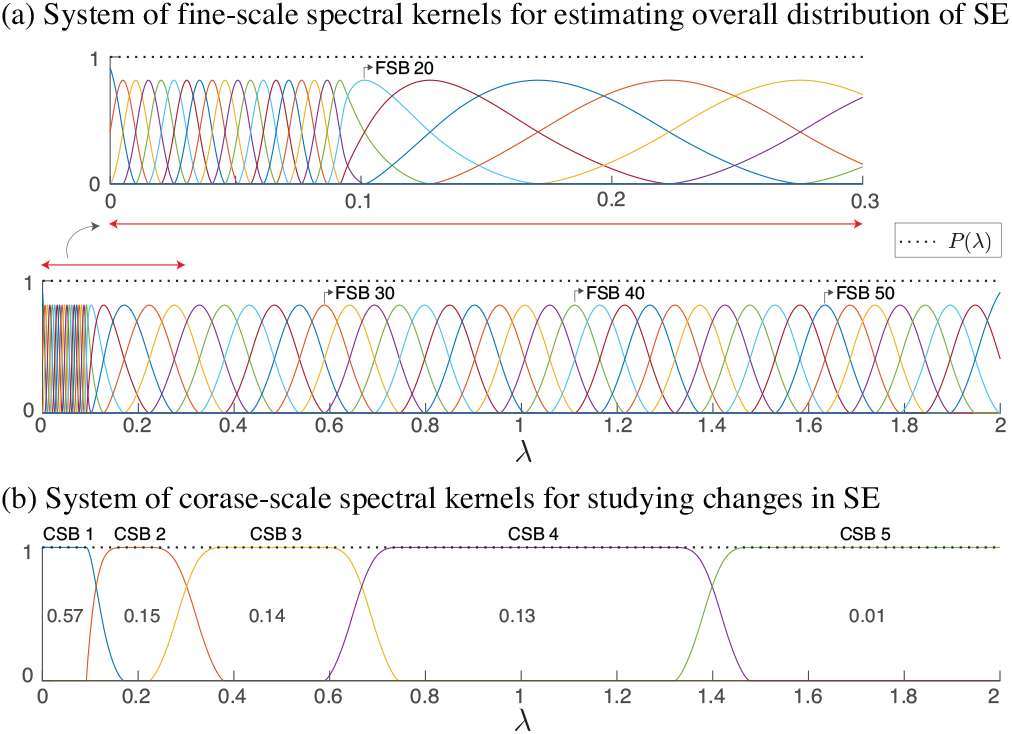
System of spectral kernels forming tight Parseval frames. In (a), kernels within the spectral range [0, 0. 1] are designed to have narrower bands as the majority of the fMRI signal energy falls in that range. In (b), values shown within the kernels represent the fraction of total SE captured within each sub-band, on average across subjects; see also Fig. 3(a).

The decomposition of **f**_*s,t*_ using each kernel 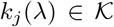 results in SE value, 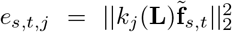, which satisfy ∑*_j_ e_s,t,j_* = 1, thanks to the normalization performed in (3) and the Parseval property of 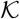. Using this measure, we characterize the ensemble distribution of energy, across *T* time frames, for a given subject as

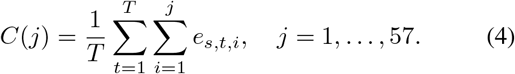

In order to reduce the dimensionality of the spectral representation, we studied variations in the SE content using a coarser set of five spectral kernels as shown in Fig. 1(b), denoted 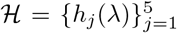, which also form a Parseval frame. Using 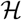, we obtained five SE values for each signal 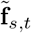 as

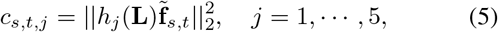

which satisfy ∑*_j_ c_s,t,j_* = 1. The span of the coarse-scale spectral bands (CSB) was determined as follows. The first CSB was set to cover approximately the lower 5% of the spectrum, i.e., λ ∈ [0,0.1], to be consistent with previous results on spectral characterization of fMRI data on cortical graphs [18]. The fifth subband was defined to cover the upper approximately 1% of the ensemble SE. The mid spectral range was split into three subbands, each of which captured an approximately equal amount of SE for WM fMRI graph signals as given by (4), based on the idea presented in [20].

## 3. RESULTS

Figure 2 shows representative WM harmonics of a representative subject. The first harmonic reflects a measure of the degree at each vertex, whereas the second harmonic—the Fiedler vector [21], splits the studied hemisphere in two. WM harmonics associated to larger eigenvalues encode more varying spatial patterns relative to those associated with smaller eigenvalues. In particular, the representative WM harmonics associated with CSBs 2 and 3, λ = 0.2 and 0.5, manifest spatial structures that are reminiscent of fiber bundles orientations, whereas that associated with CSB 4, exhibits a highly variable structure. The harmonics associated to CSB 5 exhibit spatial structures that are highly variable and localized, in contrast to the lower harmonics which manifest more delocalized patterns.

**Fig. 2:**
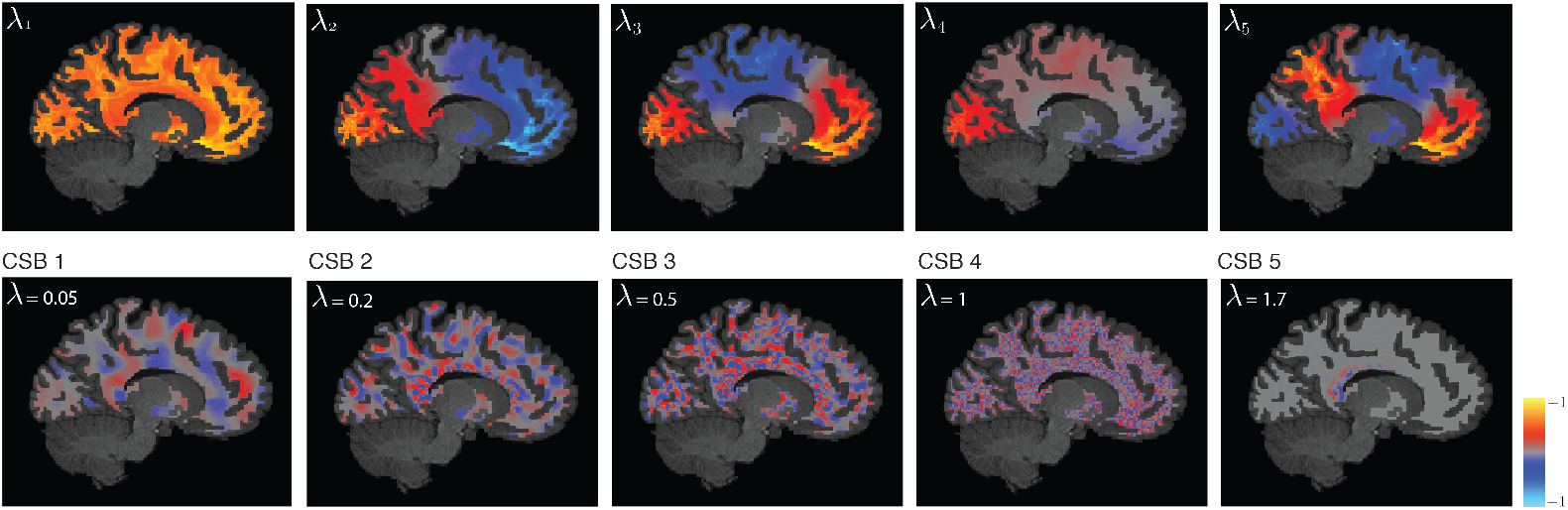
WM harmonics of a representative subject, associated with the five smallest eigenvalues (top), and eigenvalues close to the center of the five CSBs shown in Fig. 1(b). Note that the harmonics are defined in 3-D whereas a 2-D cross-section of them is displayed.

Figure 3(a) shows the distribution of SE along the spectrum, across subjects; each subject curve represents the ensemble SE aggregated across all functional frames of the Social task. More than half of the total SE is captured by WM harmonics associated to eigenvalues within the spectral range [0, 0.1], whereas less than 1% is captured by WM harmonics associated to the upper-end spectral range [1, 4, 2]. Overall, more than 90% of the ensemble signal energy is captured by eigenvectors associated to the lower half of the spectrum. WM fMRI graph signals associated to functional volumes along the five trials of the Social task were decomposed as in (5), resulting in five SE time series per CSB, per subject. The SE time series were then processed as follows: 1) smoothed with a moving average filter of length five frames (3.8 seconds), 2) de-meaned to have zero mean, 3) normalized to have a 0 onset, 4) fitted to a polynomial of order four—the choice of 4th degree was to enable fitting a curve that potentially mimics the WM HRF response, in particular, an undershoot and an overshoot, and 5) averaged across the trials for the condition, resulting in a single ensemble polynomial. A global ensemble curve was then obtained for each CSB by averaging the subject-specific ensemble curves across the 95 subjects.

**Fig. 3:**
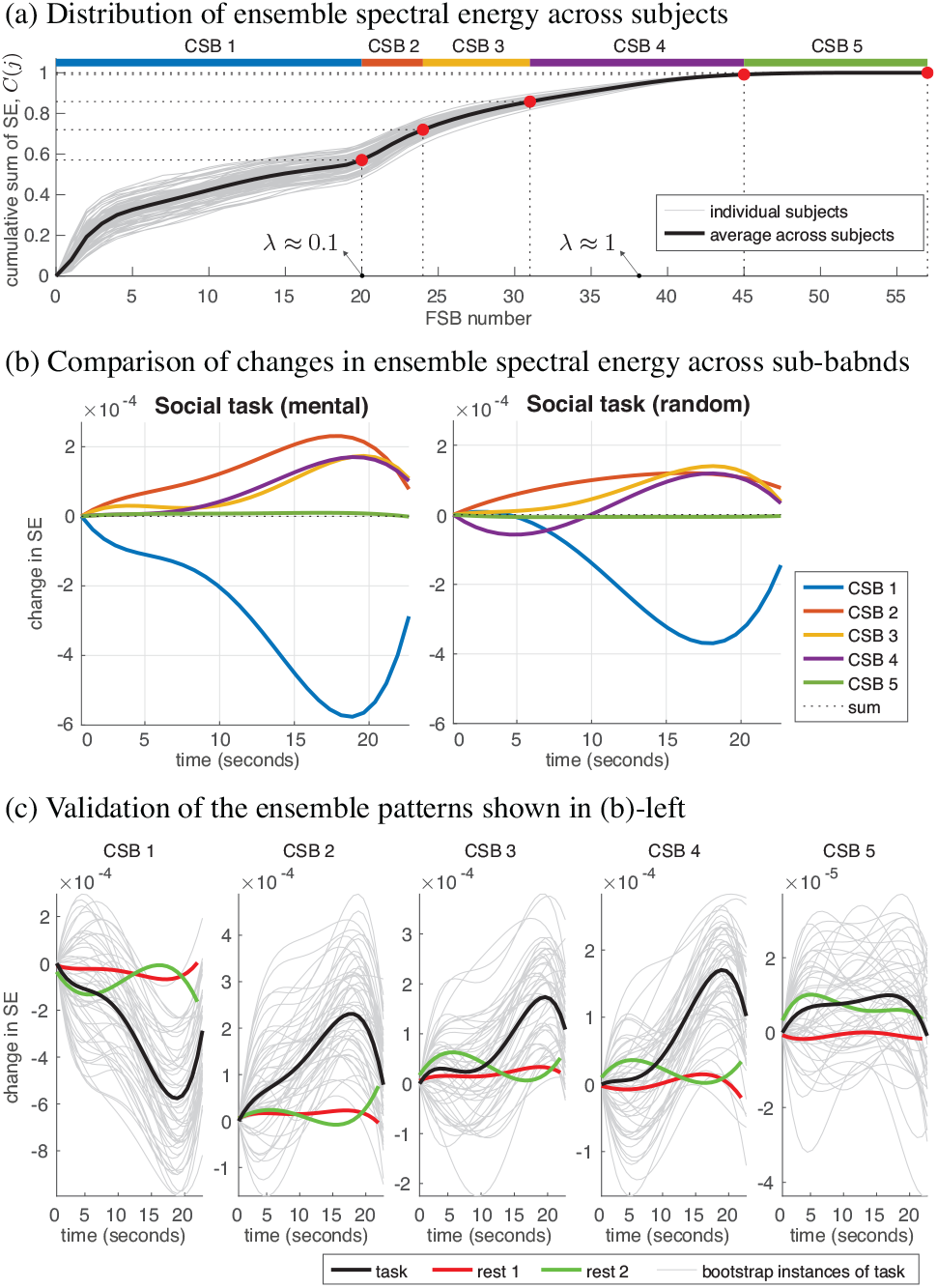
Characterization of spectral energy content of WM fMRI data using diffusion-informed WM harmonics.

Fig. 3(b) compares changes in the ensemble SE content across the CSBs in the two experimental conditions of the Social task. The SE content in CSB 1 drops during the course of the task, whereas it shows an overall increasing pattern in CSBs 2 to 4 and shows negligible variation in CSB 5. The interplay between the SE contents across the CSBs manifest subtle variations between the two experimental conditions, with the mental-condition showing a greater drop at CSB 1 relative to the random-condition, suggesting the greater engagement of more finely resolved spatial patterns during the former condition, and thus, reflecting WM spatial dynamics of varying nature underlying the two conditions. Moreover, the extent of change in the spectral content across the CSBs notably increases 10 seconds post stimulus, an observation that can be potentially linked to the delayed manifestation of HRF peaks in WM, which have also been shown to appear 10 seconds post stimulus in multiple WM fiber bundles [22, 7].

Given the low amplitudes of modulation, we performed two validations to verify the reliability of the observed patterns in Fig. 3(b). Firstly, we performed bootstrapping to see how replicable each observed pattern is, by randomly selecting 20 subjects out of the pool of 95 subjects and computing an ensemble curve, repeated 50 times; results shown in Fig. 3(c). The resulting curves reflect the replicability of the observed ensemble patterns in Fig. 3(b), which are replicated in Fig. 3(c) as black curves. Secondly, to verify that the observed patterns are related to the underlying functional task, ensemble curves over 95 subjects were computed on random segments of the subjects’ two resting-state acquisitions. Resting-state ensemble curves show substantially lower-amplitude variations compared to task curves, and furthermore, do not manifest a clear decrease/increase in SE across time, reflecting the greater stability of the underlying spatial patterns in the resting-state data relative to task data.

## 4. CONCLUSIONS AND OUTLOOK

From a broad perspective, our results show how the spatial dynamics of WM fMRI data alter under functional loading, demonstrating an interplay between contributions from slowly varying and highly varying WM harmonic, corroborating similar findings on region-based gray matter fMRI graph signals defined on the connectome [23, 24]. Methods presented in this work can find application in multiple scenarios. The quantification of changes in WM BOLD signal has been suggested as a marker for detecting cognitive decline [25], but given the low amplitudes of WM HRF [22], we anticipate changes in SE at different CSBs to be found as an alternative, more sensitive, identifier of subtle changes in the signal. Moreover, the manifestation of a clear modulation of SE at the different CSBs suggest the potential benefit of deriving white matter functional connectivity matrices [26] from fMRI data that are spatially filtered to retain contributions from a given CSB—suppressing contributions from the other CSBs, rather than smoothing the data using isotropic Gaussian filters, which deteriorate the inherent diffusion-dependent spatial structure in the data. Lastly, the decomposition of WM fMRI data at multiple CSBs can be leveraged, for example, to implement multi-scale spatial denoising of the data [27], to derive WM signatures of consciousness [28, 29], or to train models for predicting disease [30].

## 5. ACKNOWLEDGMENTS

This work, and HB, were supported by the Swedish Research Council (2018-06689), and in part by the Royal Physiographic Society of Lund, the Thorsten and Elsa Segerfalk Foundation, the Hans Werthén Foundation and the Sweden-America Foundation. IA was supported by the BrightFocus Foundation (A2016172S) and NIH grants K01DK101631 and R56AG068261. DA and AE were supported by the Swedish Research Council (2017-04889), the ITEA3 / VINNOVA funded project Intelligence based iMprovement of Personalized treatment And Clinical workflow supporT (IM-PACT), and the Center for Industrial Information Technology (CENIIT) at Linköping University. CFW was supported by NIH grants P41EB015902 and R01MH074794. The authors certify that they have no conflict of interest to report in regards to the subject matter discussed in this paper.

## 6. COMPLIANCE WITH ETHICAL STANDARDS

The present research study was conducted retrospectively using human subject data made available in open access by the HCP. The HCP study was approved by the Washington University Institutional Review Board and informed consent was obtained from all subjects. Ethical approval for analyzing the openly available data is not required according to our local ethics committee.

